# A Bivalent Chromatin Controls Timing of Expression of Camalexin Biosynthesis Genes in Response to a Pathogen Signal in Arabidopsis

**DOI:** 10.1101/2021.05.05.442749

**Authors:** Kangmei Zhao, Deze Kong, Benjamin Jin, Christina D. Smolke, Seung Y. Rhee

## Abstract

Temporal dynamics of gene expression underpins the responses to internal and environmental stimuli. In eukaryotes, regulation of gene induction includes changing chromatin states at the target genes, and recruitment of the transcriptional machinery that includes transcription factors. As one of the most potent specialized metabolites in *Arabidopsis thaliana*, camalexin can be rapidly induced by bacterial and fungal infections. Though several transcription factors controlling camalexin biosynthesis genes have been characterized, it remains unknown how the rapid activation of genes in this pathway upon a pathogen signal happens. By combining publicly available epigenomic data with *in vivo* chromatin mapping, we found that camalexin biosynthesis genes are marked with two epigenetic modifications with opposite effects on gene expression, H3K27me3 (repression) and H3K18ac (activation), to form a bivalent chromatin. Mutants with reduced H3K27m3 or H3K18ac showed that both modifications were required to determine the timing of gene expression and metabolite accumulation at an early stage of the stress response. Our study indicates that this type of bivalent chromatin, which we name a kairostat, plays an important role controlling the timely induction of gene expression upon stimuli in plants.

## Introduction

Plants produce many specialized metabolites that are essential for responding to environmental cues, the expression of which requires tight temporal control ^1, 2, 3^. One of the most prominent specialized metabolites from *Arabidopsis thaliana*, a model plant, is camalexin. Camalexin is an indole alkaloid derived from tryptophan and can be induced rapidly by various biotic and abiotic stress stimuli in Arabidopsis^4^. Essential enzymes required to catalyze reactions in this pathway belong to the Cytochrome P450 family, including *CYP79B2/3*, *CYP71A12/13*, and *Phytoalexin Deficient 3 PAD3^5, 6^*. The expression of these enzymatic genes can be rapidly activated by pathogen elicitors, such as flagellin 22 and oligogalacturonides^7^. Transcription factors and protein kinases that regulate the expression of camalexin biosynthesis genes have been identified^8, 9, 10, 11^. For example, MYB122 promotes camalexin biosynthesis under *P. syringae* infection^9^. WRKY33 functions as an activator and directly binds to the promoters of camalexin biosynthesis genes^11^. CALCIUM-DEPENDENT PROTEIN KINASE5 (CPK5) and CPK6 can phosphorylate the Thr-229 residue of WRKY33 to enhance its binding ability^10^. The accessibility of target gene regions to transcription factors is determined by the dynamics of chromatin states in eukaryotic cells^12^ and epigenetic modifications play important roles in determining gene induction through influencing nucleosome accessibility^13^. Despite the identification of transcription factors affecting camalexin biosynthesis, it remains unknown how the rapid activation of genes associated with this pathway upon a pathogen signal is enabled and what roles epigenetic modifications and chromatin conformation have in this process.

Epigenetic modifications constitute various covalent decoration of chemical groups to histones and DNA, which are associated with promoting or repressing gene expression^12, 14, 15^. For example, trimethylation of lysine 27 of histone 3 (H3K27me3), established by the Polycomb Repressive Complex 2 (PRC2), is associated with repressing gene expression^16^. H3K27me3 represses gene expression by increasing chromatin condensation and limiting the recruitment of the transcriptional machinery^17^. Trimethylation of lysine 4 of histone 3 (H3K4me3), is marked at actively transcribed genes^18^, which activates gene expression by recruiting initiation factors to the promoters of target genes^19^. Epigenetic marks that play opposite roles on gene expression can colocalize at the same gene regions to form bivalent chromatin^20, 21^. Bivalent chromatin was initially observed in embryonic stem cells and formed by H3K27me3 and H3K4me3^20, 22, 23^. In plants, the H3K27m3-H3K4me3 bivalent chromatin has been reported in *FLOWERING LOCUS C* (*FLC*)*^24^* in *Arabidopsis thaliana* and several thousand genes expressed in potato tuber^25^. In addition, H3K18ac was observed to positively correlate with H3K27me3 based on the Arabidopsis chromatin state analysis and the colocalization of these two marks were tested at a few loci, including *Golden2-like* 1 (*GLK1*), *Nitrate Reductase 2* (*NIA2*), and *Pathogen and Circadian Controlled 1* (*PCC1*)^26^. Recently, a novel bivalent chromatin state formed by H3K27me3 and H3K4me1 was identified in *Brassica napus^27^*. Therefore it appears that bivalent chromatin can be formed by different pairs of epigenetic marks. The function of bivalent chromatin has been proposed to poise the expression of developmental genes in stem cells for rapid activation^20^. However, this hypothesis has not been tested *in vivo*. Despite the broad occurrence of bivalent chromatins, their functional roles on gene expression control remains to be elucidated in any organism.

In this study, we identified epigenetic modifications associated with metabolism by integrating epigenomic profiles with a genome-wide metabolic network. The results showed that two epigenetic marks playing opposite roles on gene expression, H3K27me3 and H3K18ac, were enriched at camalexin biosynthesis and another 12 pathways associated with specialized metabolism. Focusing on the camalexin biosynthesis pathway in this study, sequential ChIP-PCR confirmed that these two marks were colocalized *in vivo* and formed bivalent chromatin. Mutants defective in H3K27m3 and H3K18ac showed that both modifications were required to determine the timely induction of gene expression and metabolite accumulation under FLG22 treatment. This study revealed novel insights on the epigenetic regulation of camalexin biosynthesis pathway and the function of bivalent chromatin on determining gene expression.

## Results

Camalexin biosynthesis pathway is one of the best characterized pathways associated with specialized metabolism in Arabidopsis^4, 8^. We identified epigenetic marks associated with the camalexin biosynthesis pathway using a data-driven approach by integrating publicly available epigenomic profiles with Arabidopsis genome-scale metabolic network. The 16 high-resolution epigenomic profiles including histone variants, DNA methylation, and histone modifications, were generated from seedlings grown under similar conditions^26, 28, 29^. To map the epigenetic marks on metabolic genes, pathways, and domains, we used the genome-wide functional annotations of metabolism we generated previously^30^. We then asked whether certain epigenetic modifications were associated with specialized metabolic genes compared to other domains of metabolism. Enrichment analysis revealed distinct patterns with enrichment of a repression mark H3K27me3 and an activation mark H3K18ac for specialized metabolic domain (Fig. 1A). Compared to all the genes marked by these two modifications, genes involved in specialized metabolism were more likely to be associated with both marks than expected by chance in the genome (hypergeometric test, p value = 0.003, fold change = 4.8) (Fig. 1B, Fig S1). To understand how prevalent these observed epigenetic modification patterns were for specialized metabolic pathways, we performed pathway-level enrichment analysis and found that 23 pathways were enriched with both H3K18ac and H3K27me3, including camalexin biosynthesis pathway (Fig. S2, Table S1).

**Fig. 1.**
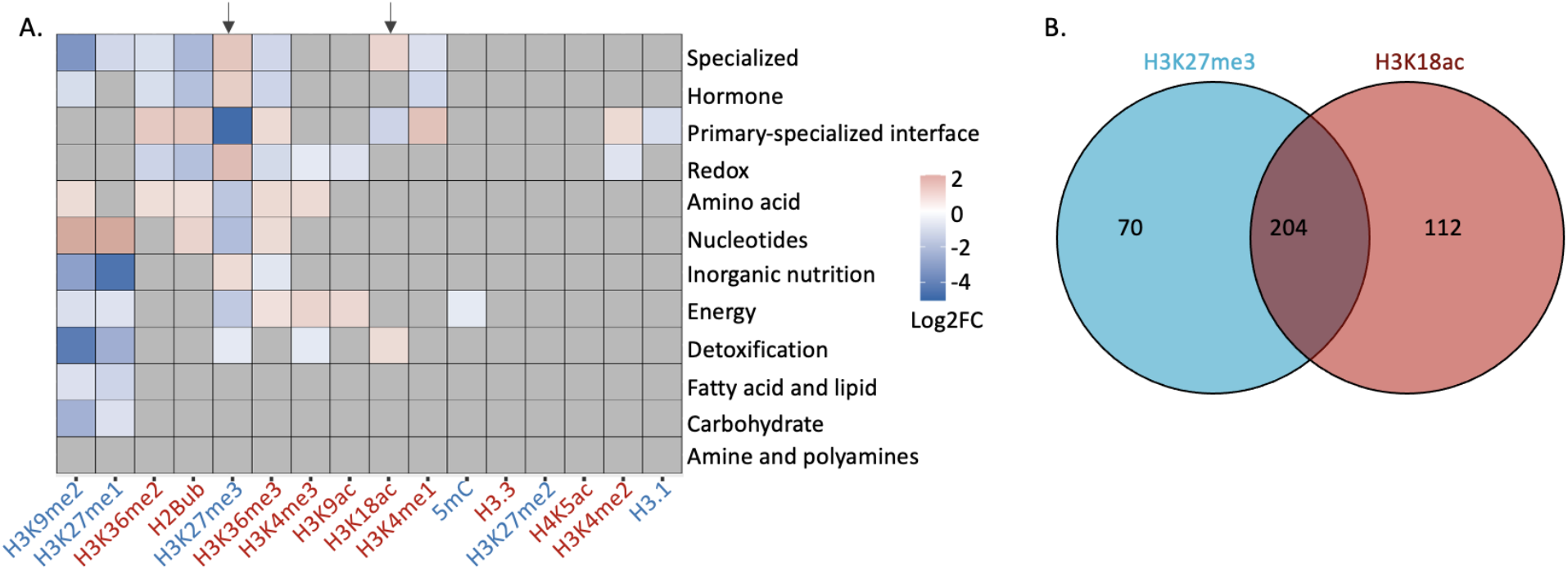
Patterns of epigenetic modification across metabolism. A. Enrichment analysis shows differential epigenetic modification patterns across metabolic domains. The heatmap represents log2 fold change (Log2FC) of enrichment or depletion of an epigenetic mark relative to total metabolic genes. Metabolic genes only mapped to unique domains were included in this analysis. Significant enrichment or depletion is based on Fisher’s exact test, p value < 0.05 and fold change > 1.5. Gray cells represent no significant change. B. The co-occurrence of H3K27me3 and H3K18ac on specialized metabolic genes.

Based on this information, we developed two mutually non-exclusive hypotheses about the role of H3K18ac and H3K27me3 on the camalexin biosynthesis gene expression: 1) H3K18ac and H3K27me3 are co-localized on the same genes involved in camalexin biosynthesis in the same cell; or 2) H3K18ac and H3K27me3 are associated with the same genes but in different cells, which may contribute to determining the cell-type specificity of this pathway. To distinguish between the two possibilities, we examined *in vivo* co-localization of the two modifications using sequential chromatin immunoprecipitation (ChIP)-qPCR, which requires a two-step, serial chromatin pull-down with antibodies against these two modifications. We focused on the three essential genes that are required to catalyze reactions Fin this pathway, including *CYP79B2*, *CYP71A13*, and *Phytoalexin Deficient 3 PAD3^5, 6^* (Fig. 2A). To assess the antibodies’ specificity and efficiency, we included various positive and negative controls in the ChIP assay and tested a transcription factor-encoding gene called *Golden-2-Like 1* (*GLK*) that was previously observed to be associated by both H3K27me3 and H3K18ac^26^. All three camalexin biosynthesis genes showed significantly higher signals when pulled down with both antibodies against H3K27me3 and H3K18ac than without any antibody or with the same antibody in the second pull-down (Fig. 2B). We observed comparable pull-down efficiency for *GLK*. Altering the order of the two antibodies for H3K27me3 and H3K18ac in the pull-down showed similar results (Fig. 2C). These results indicated that H3K18ac and H3K27me3 are co-localized at the camalexin biosynthesis genes *in planta* to form bivalent chromatin.

**Fig. 2.**
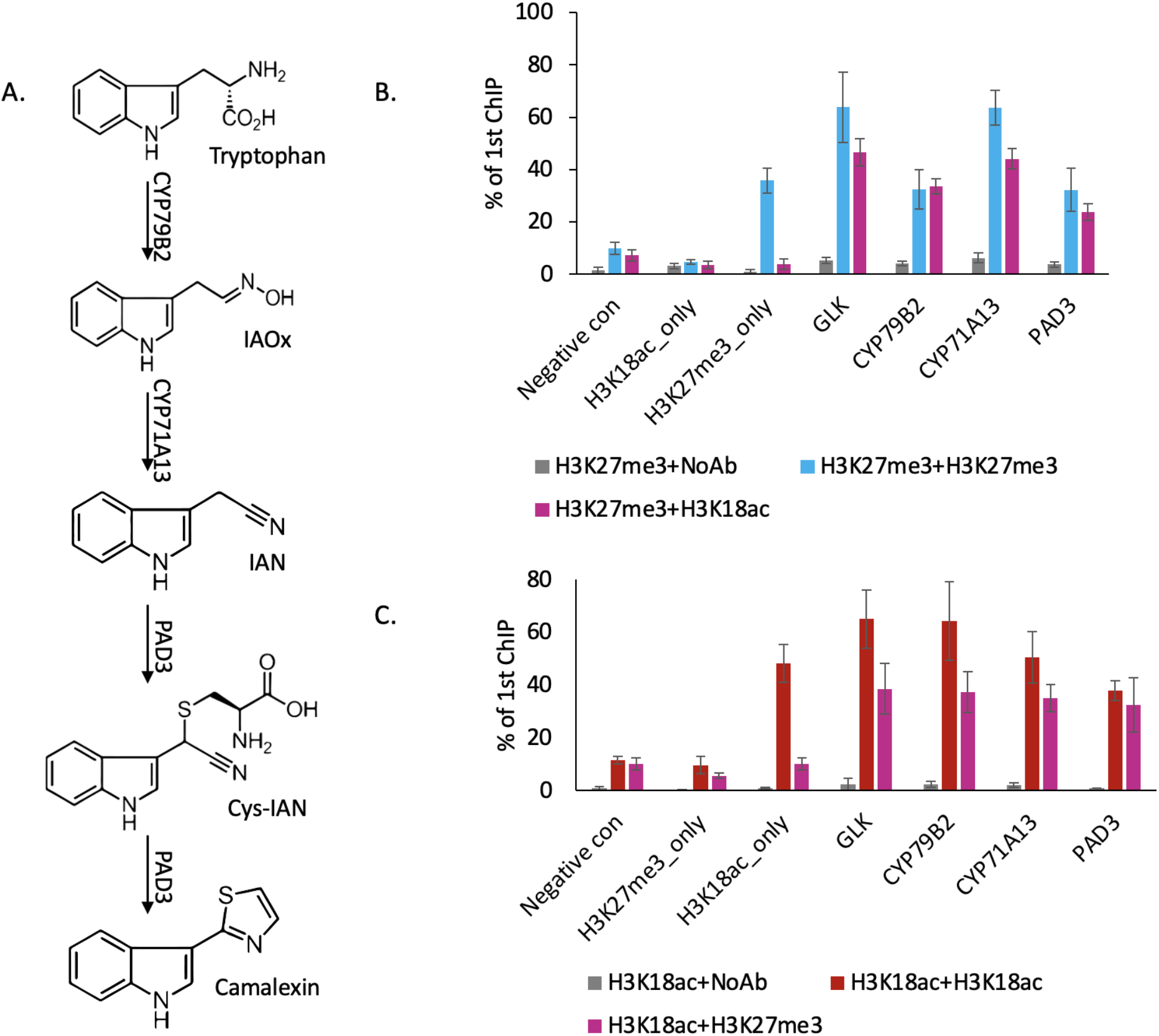
H3K27me3 and H3K18ac are co-localized on camalexin biosynthesis genes *in planta* to form bivalent chromatin. A. Camalexin biosynthesis pathway and the minimum set of genes that can produce camalexin. B-C. Sequential ChIP-qPCR confirms the co-localization of H3K27me3 and H3K18ac. Altering the order of antibodies against H3K27me3 or H3K18ac in two rounds of pull-downs shows similar results in sequential ChIP-qPCR. Negative control represents a genomic region that has very low abundance for H3K27me3 and H3K18ac. H3K27me3_only represents a genomic region that has a high abundance of H3K27me3 and very low abundance of H3K18ac. H3K18ac_only represents a genomic region that has a very high abundance of H3K18ac and very low abundance of H3K27me3. NoAb represents no antibody in the second pull-down. Error bars represent standard deviation of data from three biological replicates. The experiments were performed twice with different plant samples and revealed similar results.

The biological function of bivalent chromatin has long been hypothesized to poise gene expression for rapid activation upon signaling. To date, however, no direct evidence is available to test this hypothesis in a whole organism context. To understand the effect of bivalent chromatin on gene expression, we examined the transcriptional kinetics of camalexin biosynthesis genes under FLG22 induction in wild type and mutant lines that have defective deposition of H3K27me3 or H3K18ac. The selected lines included H3K27me3 defective mutants *curly leaf 28* (*clf28*) and chromatin remodeler *pickle (pkl-1)^16^*, and H3K18ac defective mutants *increased DNA methylation 1* and *2* (*idm1* and *idm2*)^31, 32^. In wild type plants, *CYP71A13* and *PAD3* were significantly induced within 30 minutes after FLG22 treatment and *CYP79B2* was significantly induced within 1h after the treatment (Fig. 3A to C, Fig S3, Table S5). In mutants with reduced deposition of H3K27me3 or H3K18ac, the induction patterns of the camalexin biosynthetic genes were disrupted. For *pkl-1* and *clf28* (reduced H3K27me3 marks), all three camalexin biosynthesis genes showed a significant induction of expression much faster than the wild type, within 5 minutes of FLG22 treatment, and the degree of induction was much higher than that in wild type plants at 6h of the treatment (Fig. 3A to C, Fig S3, Table S5). In contrast, in the lines with reduced H3K18ac marks, *idm1* and *idm2*, genes were induced much later than the wild type. *CYP71A13* and *PAD3* were significantly induced within 1h after the treatment and *CYP79B2* was induced within 3h after the treatment (Fig. 3A to C, Fig S3, Table S5).

**Fig 3.**
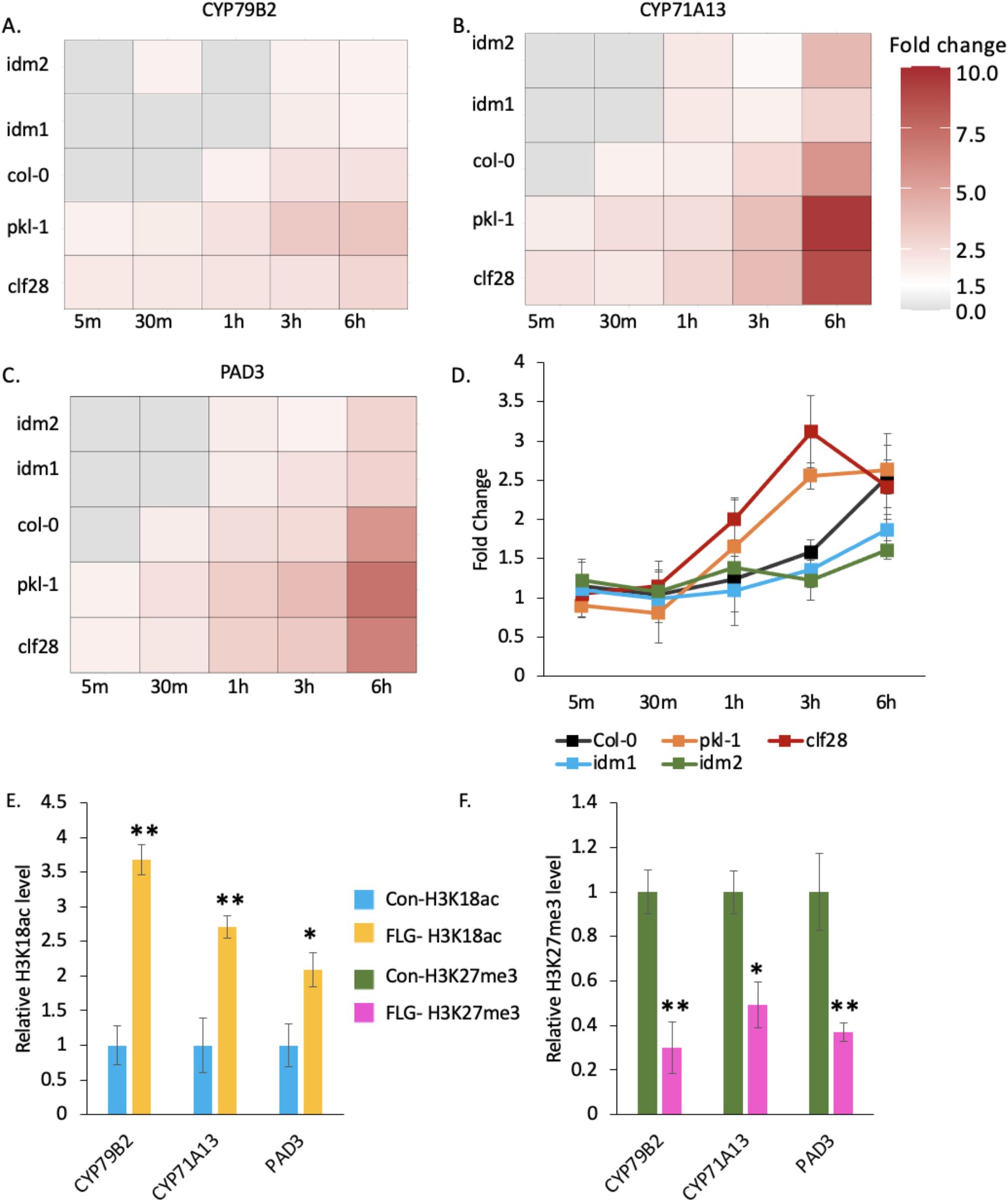
H3K27me3-H3K18ac bivalent chromatin controls the timing of gene induction and camalexin accumulation upon a pathogen signal. A-C. Expression of the three major genes in camalexin biosynthesis pathway in response to FLG22 in the wild type (Col-0), H3K27me3-defective mutants *clf28* and *pkl-1*, and H3K18ac-defective mutants *idm1* and *idm2*. ACT2 was used as a reference gene. Fold change represents the induction level of camalexin biosynthesis genes in each genotype under FLG22 treatment relative to their corresponding untreated controls at each time point. Data used to generate this heatmap and standard deviation of the fold change are summarized in Table S5. D. Camalexin accumulation in the wild type and mutants in response to FLG22. Camalexin content at each time point is reported in Table S6. Error bars represent standard deviation of data from three biological replicates per experiment. The experiments were performed twice with different plant samples and showed similar results. E-F. Abundance of epigenetic modifications at the genomic regions of camalexin biosynthesis genes with or without FLG22 treatment. Con-H3K18ac = chromatin complexes were extracted from water treated samples (Col-0) and pulled-down by the antibody against H3K18ac. FLG-H3K18ac = chromatin complexes were extracted from FLG22 treated samples (Col-0) and pulled-down by the antibody against H3K18ac. Con-H3K27me3 = chromatin complexes were extracted from water treated samples (Col-0) and pulled-down by the antibody against H3K27me3. FLG-H3K27me3 = chromatin complexes were extracted from FLG22 treated samples (Col-0) and pulled-down by the antibody against H3K27me3. Relative abundance of each modification was calculated by normalizing first to input, then to control plants with mock treatment. Two-week old Arabidopsis seedlings of Col-0 were treated with 1μM FLG22 or deionized water in these experiments. Stressed and control samples were collected 30 min after the treatment. Error bars represent standard deviation of data from three biological replicates per experiment. The experiments were performed twice using different plant samples and showed similar results. * and ** represents p values < 0.05 and 0.01, respectively from Student’s two-tailed t test.

To test whether the changes in gene expression would lead to changes in camalexin induction, we measured camalexin content using liquid chromatography–tandem mass spectrometry (LC-MS/MS). For wild type plants, camalexin accumulated significantly 3h after FLG22 treatment and continued to increase 6h after the treatment (Fig. 3D, Table S6). Plants with reduced H3K27me3 marks, *clf28*, started to accumulate more camalexin much earlier, 1h after FLG22 treatment, continued to increase at 3h, and then stabilized 6h after FLG22 treatment. On the other hand, for mutants with reduced H3K18ac marks, *idm1* and *idm2*, camalexin accumulated significantly much later, at 6h after the treatment (Fig. 3D, Table S6). These data indicate that H3K27me3-H3K18ac bivalent chromatin is required to control the timing of expression of camalexin biosynthetic genes and camalexin production in response to FLG22.

We next asked whether the amount of H3K27me3 and H3K18ac marks on the camalexin biosynthetic genes was associated with their expression kinetics in response to FLG22. To test this hypothesis, we quantified the abundance of H3K27me3 and H3K18ac on the camalexin biosynthetic genes with or without FLG22 treatment using ChIP-qPCR. For each gene tested, primers were designed to cover a genomic region including 1kb upstream and the entire transcribed region (Table S2). In response to FLG22, the abundance of H3K18ac increased in the gene regions of *CYP71A13*, *CYP79B2*, and *PAD3* (Fig. 3E). Moreover, the abundance of H3K27me3 modification was significantly depleted in the regions of these genes in response to FLG22 (Fig. 3F). Taken together, this study reveals the association of a bivalent chromatin formed by H3K27me3 and H3K18ac on the key camalexin biosynthetic genes *in planta*, and more importantly, demonstrates the role of the bivalent chromatin for the timely induction of these genes in response to the pathogen signal FLG22.

To explore whether H3K27me3-H3K18ac bivalent chromatin is associated with the induction of other specialized metabolic genes in response to FLG22, we asked whether those genes enriched with both H3K27me3 and H3K18ac showed similar induction patterns as camalexin biosynthesis genes. We profiled the transcriptome of wild type, *pkl-1*, and *idm1* plants with or without 30 minutes of FLG22 treatment. This time point and genotypes were chosen based on the transcriptional kinetics study of camalexin biosynthesis genes. In response to FLG22, 59 specialized metabolic genes were upregulated in wild type, 80 in *pkl-1*, and 35 in *idm1* lines (Fig. S4, Table S7). Among all the induced specialized metabolic genes in *pkl*-1, 35 of them were marked by both H3K27me3 and H3K18ac, which was enriched more than expected by chance (hypergeometric test p value, 1.8E-5, fold change 1.9) (Fig S5). Among these genes, 31 showed the highest induction levels in *pkl-1* and the lowest in *idm1*, just like the camalexin biosynthesis genes (Fig4A). We further asked which pathways were enriched among these genes relative to total induced genes in *pkl-1* under FLG22 treatment and identified 13 pathways based on hypergeometric test (p value < 0.01 and log2 (fold change) > 1), including camalexin biosynthesis, glucosinolate biosynthesis, and quercetin glycoside biosynthesis among others. (Fig 4B). To determine whether these changes in gene expression are indicative of changes in the accumulation of metabolites, we performed untargeted metabolomics analysis using quadrupole Time-of-Flight (qToF)-MS in wild type, *clf28*, and *idm1* plants under control and 3h of FLG 22 treatments. *clf28* line was selected since it showed higher camalexin accumulation than *pkl-1* 3h after FLG22 induction (Fig. 3D). We identified 35 peaks that showed significantly higher accumulation in response to FLG22 treatment in *clf28* plants based on one-way ANOVA (p-value 0.05). Among them, 21 metabolites showed the highest fold change in *clf28* and the lowest in *idm1* plants, including camalexin (Fig. S6). These results suggest that the H3K27me3-H3K18ac bivalent chromatin may be required to maintain temporal induction kinetics of additional pathways.

**Fig. 4.**
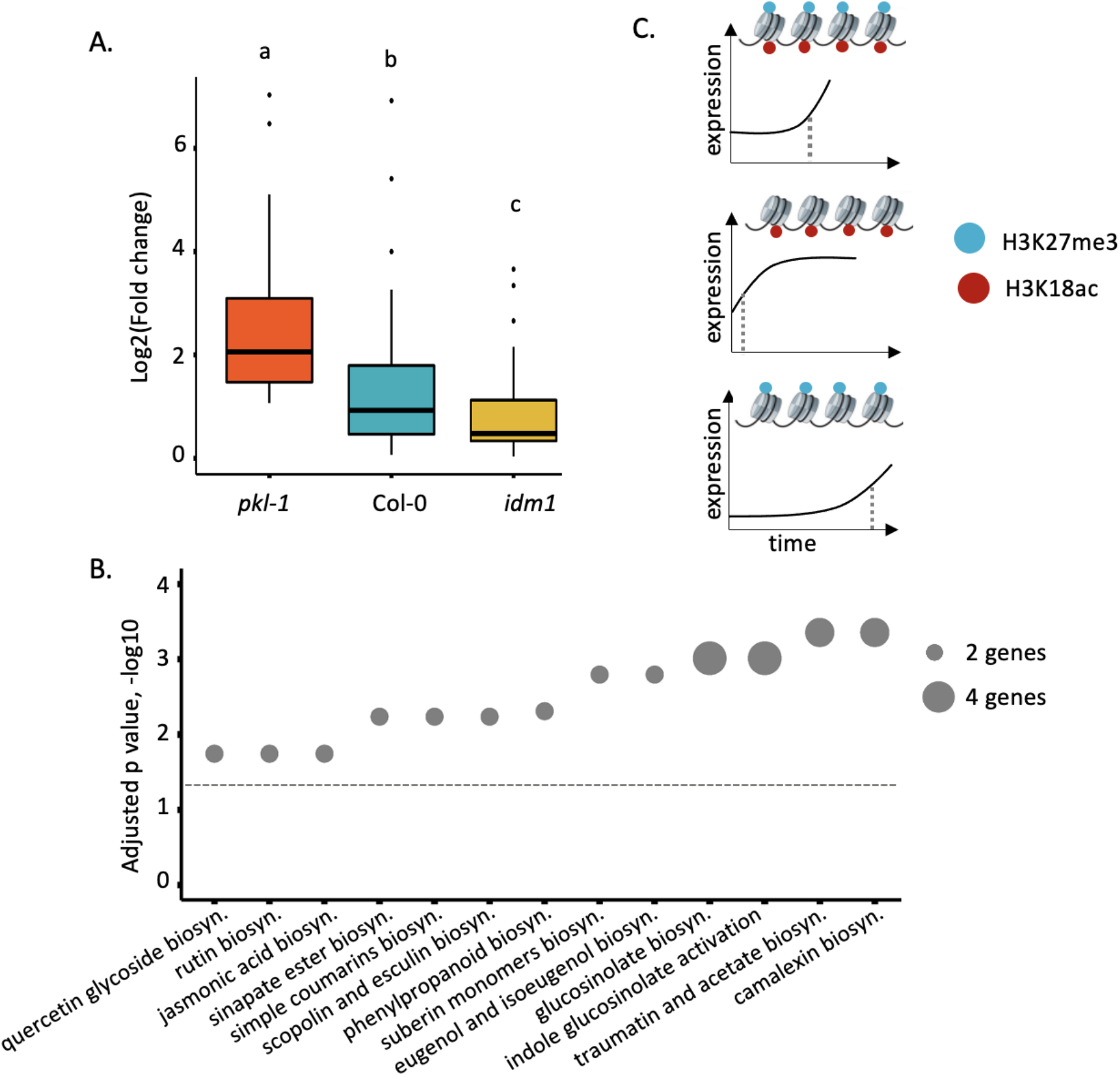
H3K27me3-H3K18ac bivalent chromatin may control additional specialized metabolic genes and pathways. A. The upregulated specialized metabolic genes in *pkl-1* marked by both H3K27me3 H3K18ac showed the highest induction level in *pkl-1* and delayed response in *idm1*. Different letters represent significant different groups of p value < 0.05 as determined by oneway ANOVA followed by post-hoc Tukey’s test. In this box plot, center lines show sample medians and box limits indicate the 25th and 75th percentiles. Whiskers extend 1.5 times the interquartile range from the 25th and 75th percentiles. Dots represent outliers. B. Enriched metabolic pathways within the induced specialized metabolic genes marked by both H3K27me3 and H3K18ac. Bubble size represents the number of induced specialized metabolic genes marked by both H3K27me3 and H3K18ac. The dashed line represents p value threshold (0.05) of the hypergeometric test. C. A model of the H3K27me3-H3K18ac bivalent chromatin that acts as a kairostat to regulate the temporal pattern of gene expression in response to external stimuli. Upon a pathogen signal, bivalent chromatin controls the timing of induction for camalexin biosynthetic genes, with H3K18ac expediting and H3K27me3 attenuating expression to hit the presumed temporal ‘sweet spot’. Dashed lines represent the time of significant gene induction.

In summary, our analysis showed that the bivalent chromatin formed by H3K18ac and H3K27me3 controls the timing of induction of camalexin biosynthesis genes and potently other specialized metabolic genes upon stress, with H3K18ac expediting and H3K27me3 attenuating expression to hit the presumed “sweet spot” (Fig. 4C). We have named this type of bivalent chromatin regulator a kairostat, inspired by the ancient Greek word ‘kairos’ meaning the right moment and ‘stat’ meaning regulating device.

## Discussion

Camalexin is the most prominent phytoalexin in Arabidopsis and genes associated with its biosynthesis are induced rapidly upon pathogen signals, such as FLG22^7, 9, 33^. Induction of gene expression requires accessibility of the chromatin and transcription factor binding to the promoters of target genes^13, 34^. Despite the identification of transcription factors controlling camalexin accumulation, it remains unknown how epigenetic modifications enable the rapid accessibility of target gene regions upon a pathogen signal. In this study, we found that H3K27me3 and H3K18ac colocalized on camalexin biosynthesis genes forming a bivalent chromatin. We functionally examined the role of H3K27me3-H3K18ac bivalent chromatin on regulating the induction of camalexin biosynthesis genes by combining publicly available epigenomics data with genetics, molecular biology, and biochemistry. In wild type (Col-0) plants, camalexin biosynthesis genes were activated at 1h of FLG22 treatment and continued to increase their expression at later time points of the treatment. This is consistent with the previous study on induction kinetics of defense pathways under FLG22 treatment^7^. In mutants with defective deposition of H3K27me3 or H3K18ac, the induction kinetics of camalexin biosynthesis genes was accelerated or delayed, respectively (Fig3). We confirmed that the changes in gene expression patterns were corroborated with changes in camalexin levels and the epigenetic marks. Together, these data indicate that H3K27me3-H3K18ac bivalent chromatin is critical to maintain the timing of induction for camalexin biosynthesis genes.

Bivalent chromatin was initially observed in embryonic stem cells with the best understood form being H3K27me3 and H3K4me3^20, 21, 22, 23^. In embryonic stem cells, lineage-specific genes involved in differentiation are associated with H3K27me3-H3K4me3 bivalent chromatin, which are expressed at low levels before differentiation^20^. The function of bivalent chromatin was hypothesized to poise developmental genes in embryonic stem cells for rapid activation upon a cell differentiation signal^20^. In plants, H3K27me3 and H3K4me3 were colocalized at *FLC* locus in Arabidopsis wild type seedlings and reducing H3K27me3 suppressed the deposition of H3K4me3 based on the comparison of H3K4me3 abundance between wild type and the *clf* mutant with defective H3K27me3 deposition^24^. In potato tuber, H3K27me3-H3K4me3 bivalent chromatin was detected at the genes associated with development and stress response under cold stress^25^. Developmental genes with the H3K27me3-H3K4me3 bivalent chromatin showed reduced expression, whereases stress response genes increased expression under cold stress^25^. Based on these results, the authors proposed that cold-induced H3K27me3-H3K4me3 bivalent chromatin may represent a chromatin state of altering expression by working with additional proteins such as LHP1 (Like Heterochromatin Protein1)^25^. In this study, we examined the role of bivalent chromatin on gene expression using mutants that are defective in making those marks and found that H3K27me3-H3K18ac bivalent chromatin is critical for the induction kinetics of camalexin biosynthesis genes upon FLG22 signal. Changes in the expression of camalexin biosynthesis genes were associated with changes in the abundance of H3K27me3 and H3K18ac (Fig3 E to F). These results indicate that H3K27me3 and H3K18ac are required to regulate the timely induction of camalexin biosynthesis genes with H3K27me3 attenuating gene expression and H3K18ac promoting gene expression.

To fully elucidate how bivalent chromatin integrates environmental cues to regulate gene expression, it would be critical to understand how bivalent chromatin is established and maintained. Specifically, it remains to be elucidated how both marks recognize common targets to establish bivalent chromatin in the genome. Mechanisms that enable histone-modification protein complexes to alter the abundance of each mark upon stress signals also remains to be elucidated. A recent study showed that NF-Y transcription factors can interact with histone modification protein REF6 to reduce H3K27me3 at *SOC1* locus^35^. It would be interesting to examine whether transcription factors that directly bind to promoters of camalexin biosynthesis genes interact with proteins that can deposit or remove both epigenetic marks forming bivalent chromatin upon stress signals. The H3K27me3-H3K18ac bivalent chromatin and its role on the camalexin biosynthesis genes described here provide the molecular handle with which to open these lines of investigations in the future. With these knowledge gaps filled, kairostats can be engineered to function as a regulatory cassette to control the expression of target genes under specific tissues and conditions. This capability will provide novel routes to re-program biological processes involved in development and stress response, which have far-reaching implications in agriculture, engineering, and medicine.

## Methods

### Annotation of Metabolic Genes, Pathways, and Domains

Arabidopsis metabolic enzymes and pathways were extracted from Plant Metabolic Network (PMN) (http://www.plantcyc.org/). Enzymes were annotated based on the Ensemble Enzyme Prediction Pipeline (E2P2) v.3.0 ^30, 36^, metabolic pathways were predicted by Pathway Tools software v.18.5^37^, which were further validated by a process called Semi-Automated Validation and Integration (SAVI) v.3.02^38^. In *A. thaliana*, 5451 genes were predicted to encode enzymes that catalyze 3585 reactions in small molecule metabolism, which were assigned to 627 pathways based on prediction and manual curation using AraCyc v.13.0^30^. These pathways were further grouped into 13 metabolic domains based on the types of metabolites that they potentially synthesize^30^. The metabolic domain(s) assigned to a given pathway were transferred to the reactions, EC numbers, and proteins/genes associated with that pathway. To improve the statistical power, we included only those pathways that contained more than ten genes when we performed epigenetic modification enrichment analysis on individual pathways in specialized metabolism^39^.

### Epigenomic Enrichment Analysis

The 16 epigenomic profiles were downloaded from Wang et al. 2015, where previously published ChIP-seq and ChIP-chip data for all epigenetic marks were re-mapped to *Arabidopsis thaliana* genome TAIR v.10^28^. Briefly, for each ChIP-seq profile, the signals were binned and ranked in the genome^28^. We used the ranked epigenetic modification signals in the following analyses.

To identify the predominant epigenetic modifications associated with metabolic domains and pathways, we first mapped the ranked epigenetic modification signals to each gene, including the transcribed region, 1kb upstream, and 500bp downstream regions. Then, we calculated the average epigenetic modification density for each domain or pathway by taking the sum of the ranked modification signals observed within all the genes in that domain or pathway and normalizing it to the total length of all genes. To identify enriched marks within metabolic domains or pathways, we compared the density of each epigenetic modification within that domain or pathway to the background signal. We defined the background signal as the average density of each epigenetic modification by taking the sum of the epigenetic modification signals observed within total metabolic gene regions and normalizing it to the length of these genes. The statistical significance of enrichment or depletion of epigenetic marks per domain or pathway was determined by fold change of at least 1.5 and Fisher’s exact test p value, followed by a post-hoc adjustment using FDR with the threshold of 0.01. The heatmaps were generated with gplots v.3^40^ and ggplot2 v.3.1^41^ packages in RStudio.

### Plant Materials and Growth Conditions

Seeds of *Arabidopsis thaliana* accession Columbia (Col-0) (wild type) and previously characterized mutant lines *clf28* (SALK_139371), *pkl-1* (SALK_010693), *idm1* (SALK_062999), and *idm2* (SALK_138229) were used in this study. Seeds of these mutants were obtained from Arabidopsis Biological Resource Center. In all the experiments described in this study, seeds were stratified at 4°C for three days before germination and grown in 0.5X Murashige and Skoog (MS) medium containing 1% sucrose and 0.8% agar at 23°C under long days (16 h light/8 h dark) with controlled light at 100 μmol photosynthetic photons/m^2^ s (PPFD). Two-week old seedlings were harvested for the ChIP-qPCR, sequential ChIP-PCR, transcriptional kinetics, camalexin content measurement, and metabolomics experiments.

### ChIP-qPCR and Sequential ChIP-PCR

Two-week old Arabidopsis seedlings (Col-0) were used in ChIP and sequential ChIP experiments. To assure enough chromatin for immunoprecipitation, 200 seedlings (about 2g) were pooled for each ChIP reaction. Plant tissues were fixed with crosslinking buffer containing 1% formaldehyde for 20min by vacuum infiltration at room temperature and quenched with glycine (final concentration 100 mM) for 5min. After grinding, the cross-linked chromatin was isolated using ice-cold nuclear lysis buffer (50 mM HEPES at pH 7.5, 150 mM NaCl, 1 mM EDTA, 1% Triton X-100, 0.1% Na deoxycholate, 0.1% SDS, and protease inhibitor) and disrupted by sonication to yield DNA fragments of 200 to 600bp^42, 43^.

To examine the abundance of H3K27me3 and H3K18ac 30 min after 1μM FLG22 treatment, ChIP-qPCR was used to quantify their abundance throughout the genomic region and 1kb upstream of the three major camalexin biosynthesis genes. After chromatin crosslinking and shearing, DNA/protein complexes were equally divided and immunoprecipitated with antibodies against H3K27me3 (Milipore, Cat No 07-449), H3K18ac (Abcam, Cat No ab1191), or IgG as antibody control (Milipore, Cat No 12-370) by referring the ChIP protocol established using Arabidopsis tissues^42, 43^. NA was purified using QIAquick PCR Purification Kit (Qiagen, Cat No 28104). Quantitative real-time PCR was performed using LightCycler^®^ 480 System (Roche) with the SensiFAST Sybr No-Rox mix (Bioline, Cat No BIO98020) from Bioline. Three biological replicates were used in each experiment. The ChIP experiments were performed twice with different Arabidopsis tissues and revealed similar results. Primers were designed to cover the 1kb upstream of the transcription start site and the entire transcribed region (Table S2).

Sequential ChIP experiments were performed to examine *in planta* co-localization of H3K27me3 and H3K18ac on camalexin biosynthesis genes. Chromatin crosslinking and shearing followed the protocol described above for ChIP-qPCR. For each ChIP/re-ChIP assay, 20 μg DNA/protein complexes were immunoprecipitated with anti-H3K27me3 (Milipore, Cat No 07-449) and anti-H3K18ac (Abcam, Cat No ab1191) antibodies using the Re-ChIP-IT kit (Active Motif, Cat No 53016). Quantitative real-time PCR was performed using LightCycler^®^ 480 System (Roche) with the SensiFAST Sybr No-Rox mix (Bioline, Cat No BIO98020) from Bioline. To represent final results after two-step chromatin immunoprecipitation, the pull-down efficiency was calculated as the percent of first ChIP using the equation: 1^st^ ChIP = 2^^(Cq_2nd ChIP - Cq_1st ChIP)^. Three biological replicates were used in each experiment. The sequential ChIP was performed twice using different Arabidopsis tissues and altering the order of antibodies in two rounds of pull-down showed similar results. Primer sequences used in this assay are provided in Table S3.

### Transcriptional Kinetics Analysis

Two-week old seedlings grown in MS agar media as described in Plant Materials and Growth Conditions were used to study transcriptional kinetics. Flagellin 22 (FLG22), a short peptide of 22 amino acids (sequence: QRLSTGSRINSAKDDAAGLQIA), was synthesized at Stanford Protein and Nucleic Acid (PAN) Facility. To induce the expression of camalexin biosynthesis genes, seedlings were sprayed with 1μM FLG22 dissolved in deionized (DI) water. Control samples were sprayed with the same volume of DI water. FLG22 and water-treated tissues were sampled at 5min, 30min, 1h, 3h, and 6h after treatment and frozen immediately in liquid nitrogen.

After grinding in liquid nitrogen, RNA was extracted using RNeasy Plant Mini Kit (Qiagen, Cat No 74904) and 2μg RNA was used for cDNA synthesis using SuperScript First-Strand Synthesis System (Thermo Fisher, Cat No 18080051). Quantitative real-time PCR was performed using LightCycler^®^ 480 System (Roche) with the SensiFAST Sybr No-Rox mix (Bioline, Cat No BIO98020) from Bioline. Primers used in this experiment are summarized in Table S4. A housekeeping gene, Actin 2 (AT3G18780), was used as the reference gene in the qPCR experiments. Three biological replicates were used in each experiment by polling 10 seedlings per replicate. The transcriptional kinetics assays were performed twice and showed similar results.

### RNA-seq Analysis

Two-week old seedlings of Col-0, *pkl-1*, and i*dm1* plants were used for RNA-seq analysis. Seedlings were treated with 1 μM FLG22 or DI water and collected 30 min after the treatment as described in the previous section. RNA was extracted using Qiagen RNeasy Plant Mini Kit (Cat No 74904). Three biological replicates were used in each experiment by polling 10 seedlings per replicate. Library preparation and sequencing were performed by NovoGene. Raw reads were mapped to Arabidopsis genome Araport 11 using STAR package v2.6 with default parameters. Normalization and identification of differentially expressed genes were conducted using the DESeq2 package v.3.12^44^ in Rstudio. Total up-regulated genes were summarized in Table S7.

### Camalexin Content Quantification Using LC-MS

Two-week old seedlings of Col-0, *pkl-1*, *clf28*, *idm1*, and i*dm2* plants were used to measure camalexin accumulation. Tissues were treated with FLG22 or DI water as described in the RNA-seq Analysis section. Metabolites were extracted using a 75% methanol solution with 0.1% formic acid. For each sample, 150 mg of ground plant tissue was dissolved in 200 μL extraction solution. The extracted samples were vortexed at room temperature for 40 min and centrifuged at 32,000 g for 10 min to pellet plant debris. The supernatant was analyzed by an Agilent 1260 Infinity Binary HPLC paired with an Agilent 6420 Triple Quadrupole LC-MS/MS, with a reversed-phase column (Agilent EclipsePlus C18, 2.1 × 50 mm, 1.8 μm), water with 0.1% formic acid as solvent A and acetonitrile with 0.1% formic acid as solvent B, at a constant flow rate of 0.4 mL/min and an injection volume of 10 μL. The following gradient was used for metabolite separation: 0-0.5 min, 10-50% B; 0.5-5.5 min, 50-98% B; 5.5-6 min, 98% B; 6.00-6.01 min, 98-10% B; 2 min post-time equilibration with 10% B. The LC eluent was subjected to MS for 0.01-6.01 min with the ESI source in positive mode, gas temperature at 350°C, gas flow rate at 11 L/min, and nebulizer pressure at 40 psi. To detect camalexin, we set the multiple reaction monitoring (MRM) ion transition to be 201.26 → 59.2, fragmentor at 135V, and collision energy at 40V. The MRM ion transitions in this work were derived from product ion scan with precursor ion set at 201.26 and the most abundant product ion was chosen for MRM transition quantification. LC-MS/MS data files were analyzed using Agilent MassHunter Workstation software v.B.06.00. Metabolite content was quantified by integrating peak area under the ion count curve. The ion counts were calibrated to camalexin chemical standard (Cat Num Sigma SML1016) and converted to measurements of titer in molar concentrations (nM). Three biological replicates were used in each experiment by polling 50 seedlings per replicate. The experiments were conducted twice with different plant samples and showed similar results.

### Metabolomics Data Collection and Analysis

Two-week old seedlings of Col-0, *clf28*, and i*dm1* plants were treated with 1 μM FLG22 or water and tissues were collected 3 h after the treatment. Metabolites were extracted using a 75% methanol solution with 0.1% formic acid. For each sample, 250 mg of ground plant tissue was dissolved in 200 μL of the extraction solution. Extracted samples were vortexed at 4°C for 45 min and centrifuged at 32,000 g for 10 min to pellet plant debris. The supernatant was filtered with 0.45 μm PTFE filters prior to metabolite analysis. Samples were analyzed by an Agilent 1260 Infinity Binary HPLC paired with an Agilent 6545 Quadrupole Time-of-Flight (qToF)-MS, with a reversed-phase column (Agilent EclipsePlus C18, 2.1 × 50 mm, 1.8 μm), water with 0.1% formic acid as solvent A, and acetonitrile with 0.1% formic acid as solvent B, at a constant flow rate of 0.4 mL/min and an injection volume of 10 μL. The following gradient was used for compound separation: 0-0.40 min, 3% B; 0.40-20.40 min, 3-97% B; 20.40-22.40 min, 97% B; 22.40-22.41 min, 97-3% B; 22.41-24.00 min, 3% B. The LC eluent was directed to the MS for 0.01-24.00 min with ESI source in positive mode, gas temperature at 350°C, gas flow rate at 11 L/min, nebulizer pressure at 35 psi, Vcap at 3500 V, fragmentor at 150 V, skimmer at 65 V, octupole 1 RF Vpp at 750 V. A mass range of 100-1700 m/z was scanned with acquisition scan rate at 1 spectra/s.

To process metabolomics data, QToF-MS profiles were converted to mzXML files using MSConvert, and untargeted metabolomics peak extraction was performed using the xcms package v.3.12^45^ in R, with a custom R script, which are available at the Smolke Laboratory GitHub page: https://github.com/smolkelab/arabidopsis_metabolomics_analysis). Among all the detected compounds, 35 of them were statistically increased in *clf28* plants 3h after 1 μM FLG22 treatment relative to the corresponding control (water-treated) samples based on one-way ANOVA. We then extracted the average abundance of these metabolites detected in Col-0 and *idm1* plants and only kept the ones that showed the highest accumulation levels in *clf2*8 and the lowest in *idm1* plants. The metabolomics experiments were performed once. Three biological replicates were used in each experiment by polling 50 seedlings per replicate.

## Data Availability

The data that support the findings of this study are available within the article and its supplementary information files. Arabidopsis metabolic genes and pathways are available at Plant Metabolic Network (https://plantcyc.org/). The RNA-sequencing data generated in this study will be deposited to the National Center for Biotechnology Information (NCBI) before publication.

## Acknowledgments

We thank Jennifer Brophy and Flavia Bossi for their critical feedback on this work. We acknowledge Jia-Ying Zhu and Yuchun Hsiao for help with ChIP-qPCR experiments and Hye-In Nam for help with growing plants.

## Author contributions

K.Z. and S.Y.R. conceived the project. K.Z. conducted computational analysis, ChIP, and transcriptional kinetics, RNA-seq experiments and analyzed camalexin accumulation. D.K. measured and quantified camalexin content from plant extracts, conducted metabolomics data collection and analysis under C.D.S.’s supervision. B.J. isolated homozygous mutants for *idm1* and *idm2*. S.Y.R. supervised this study. K.Z. and S.Y.R. wrote the manuscript and K.Z., D.K., C.D.S., and S.Y.R. edited it.

---

**Fig S1.**
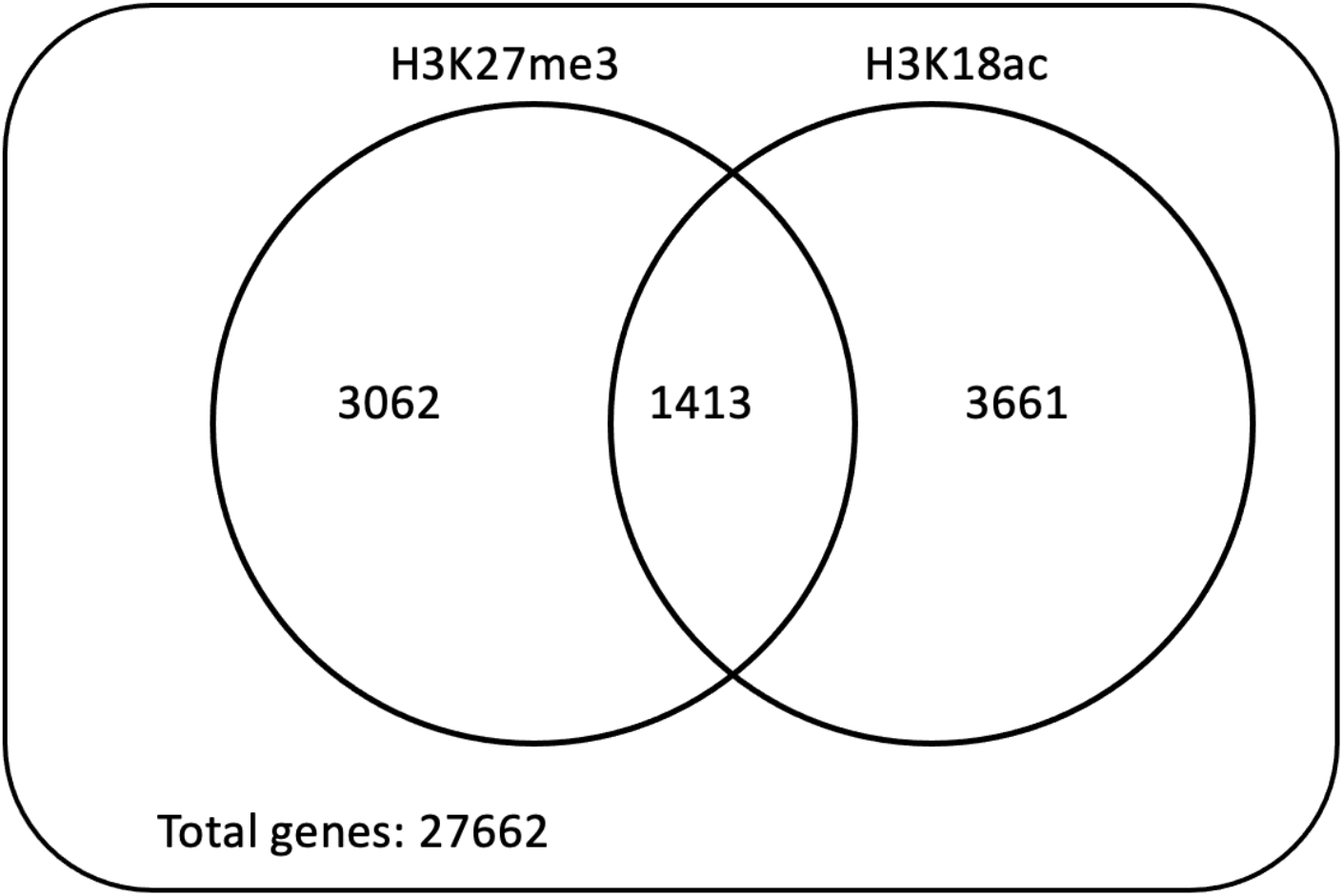
The co-occurrence of H3K27me3 and H3K18ac on total genes in the genome. 27662 represents the number of primary transcripts in Arabidopsis genome Araport 11.

**Fig S2.**
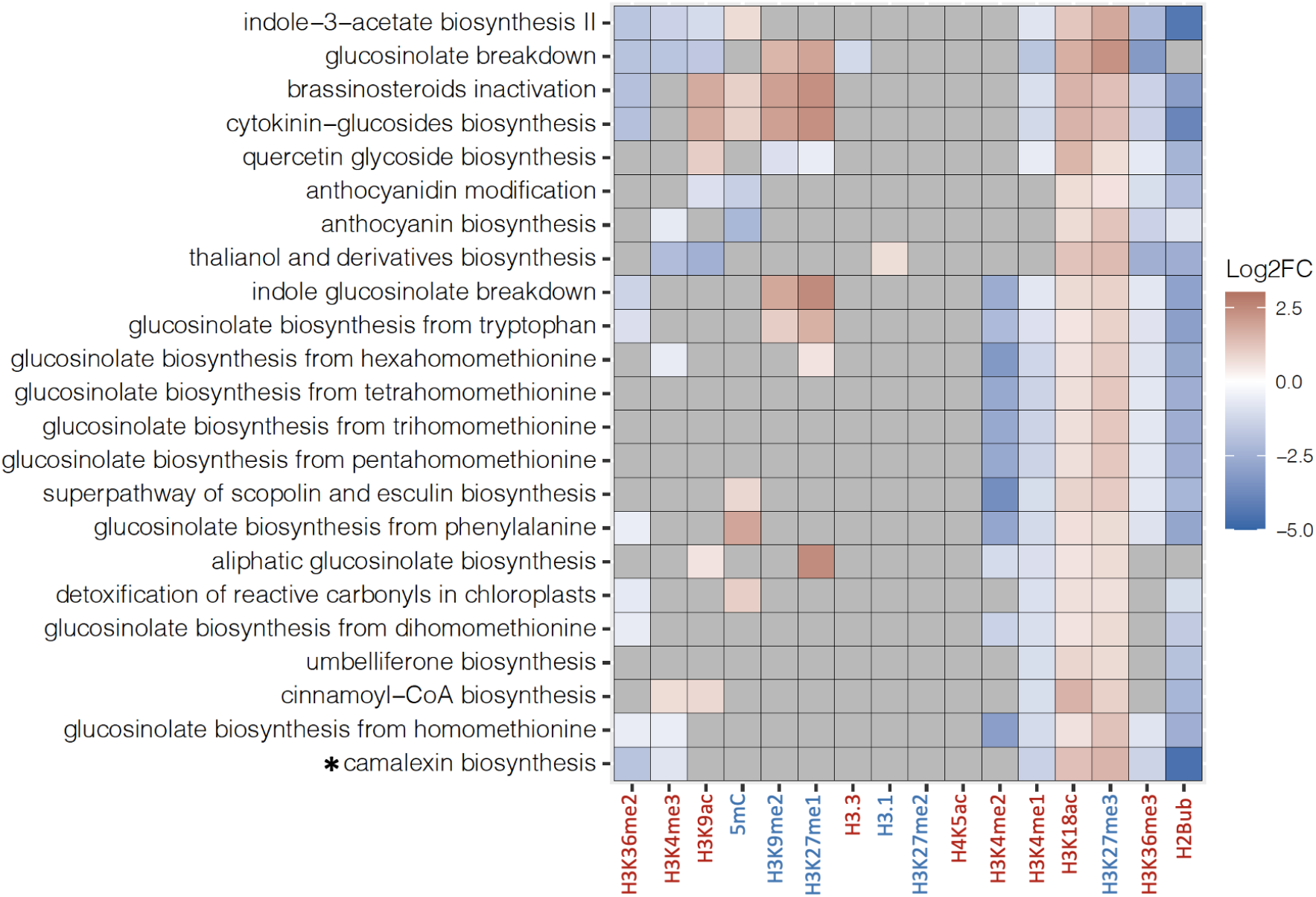
Epigenetic marks associated with pathways involved in specialized metabolism. The heatmap represents log2 fold change of enrichment or depletion of a mark relative to total metabolic genes based on Fisher’s exact test followed by a post-hoc adjustment using False Discovery Rate (FDR). Gray cells represent no-significant enrichment or depletion at FDR of 0.01 and fold change of 1.5. The heatmap was generated using hierarchical clustering using the ggplot2 package version 3.1 in RStudio. Epigenetic modifications are color-coded based on their effect on gene expression; red represents activation marks and blue represents repression marks.

**Fig S3.**
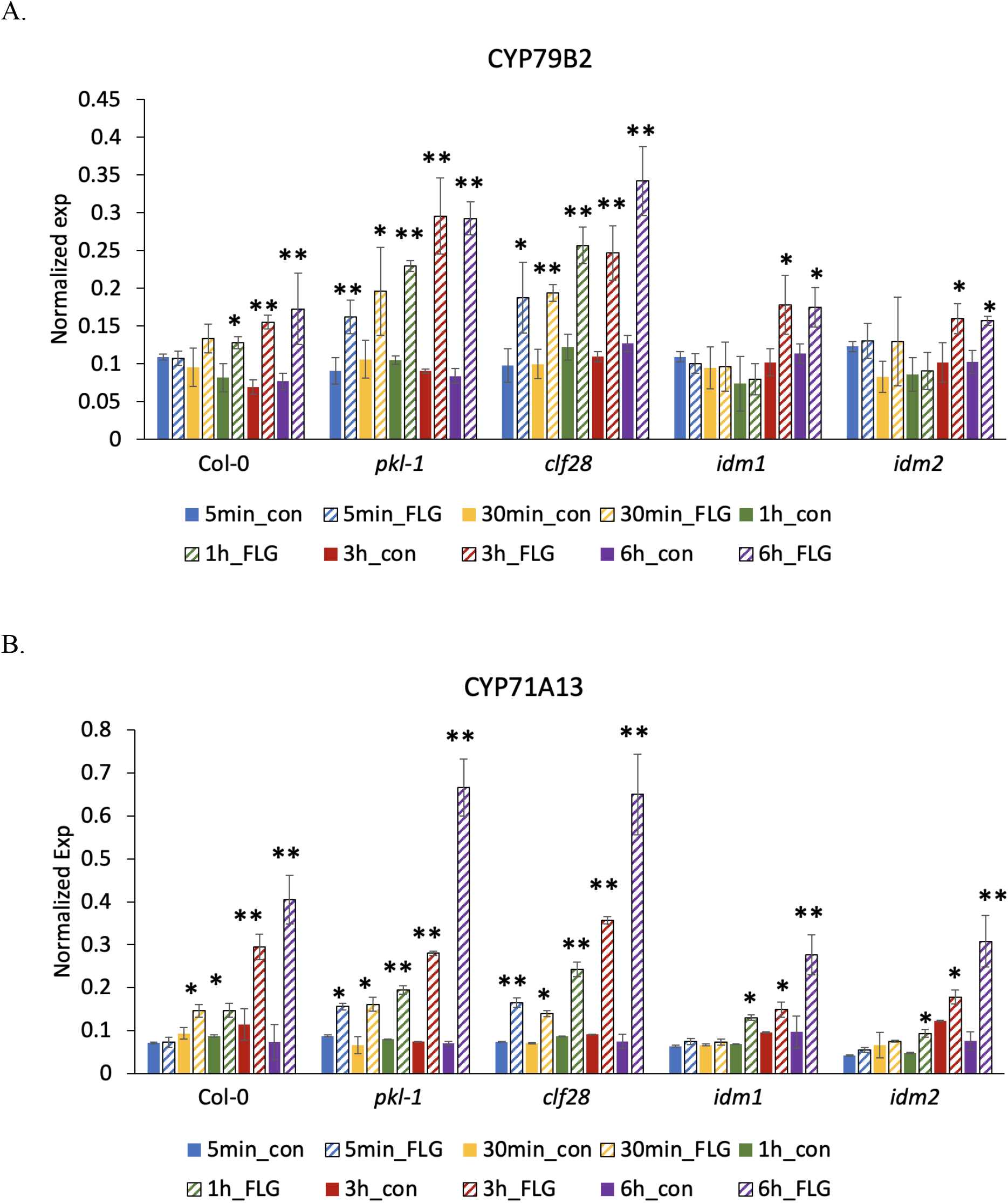
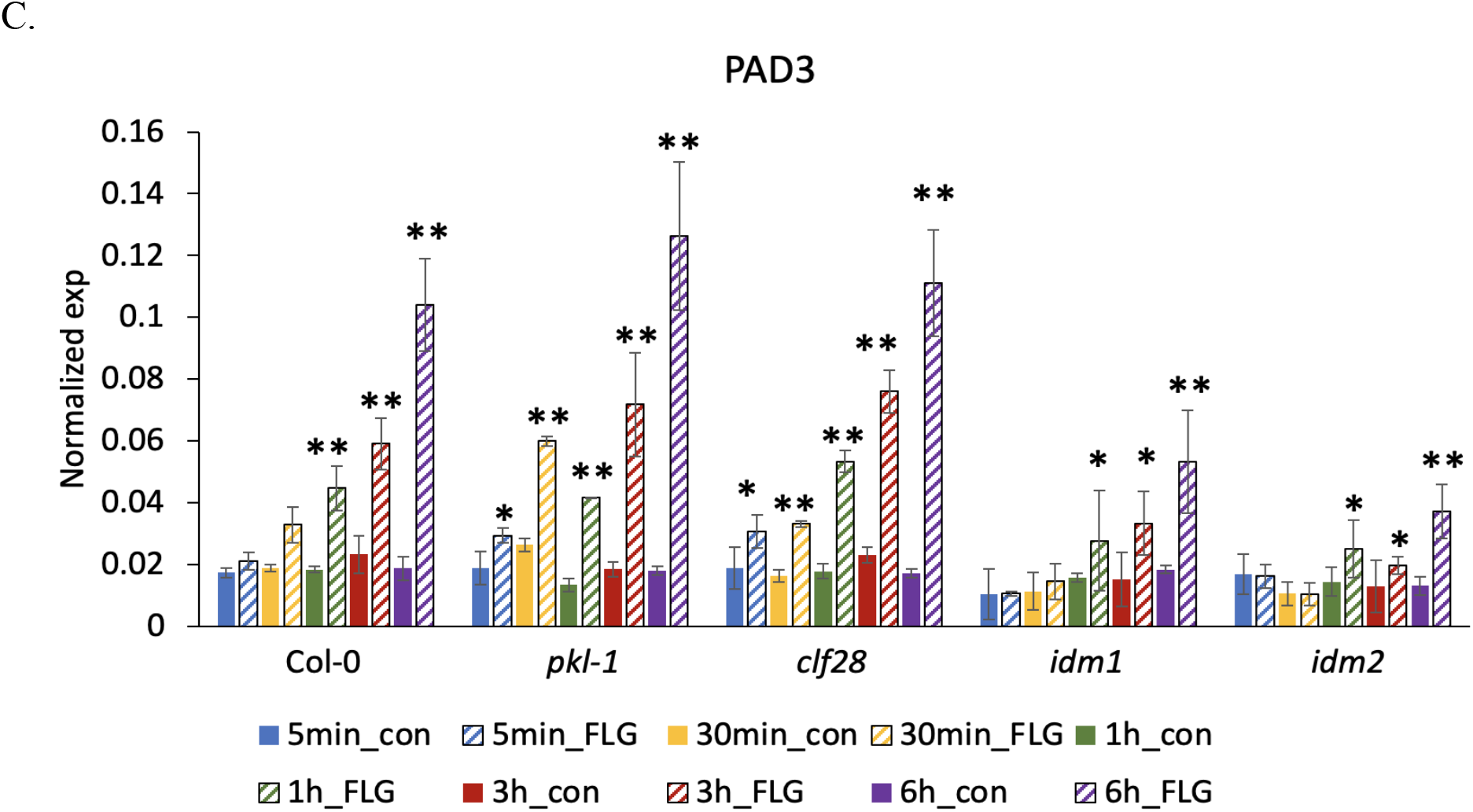
Normalized expression of camalexin biosynthesis genes upon FLG22 induction in the wild type (Col-0), H3K27me3-defective mutants *clf28* and *pkl-1*, and H3K18ac-defective mutants *idm1* and *idm2*. ACT2 was used as a reference gene. Error bars represent standard deviation of data from three biological replicates per experiment. * and ** represent Student’s two-tailed t test p value lower than 0.05 and 0.01, respectively.

**Fig S4.**
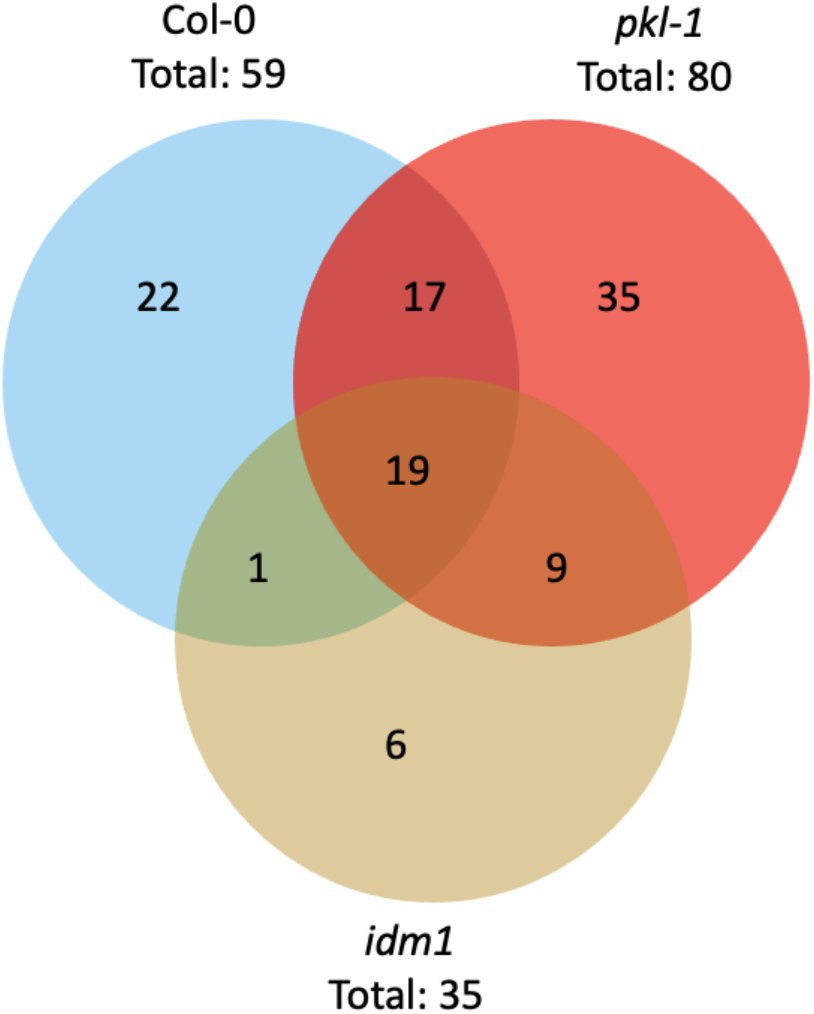
Significantly induced specialized metabolic genes in Col-0, *pkl-1*, and *idm1* by 30 min of FLG22 relative to water treatment. The criteria used to identify significantly induced genes were Log2(Fold change) > 1 and false discovery rate-adjusted p value < 0.01.

**Fig S5.**
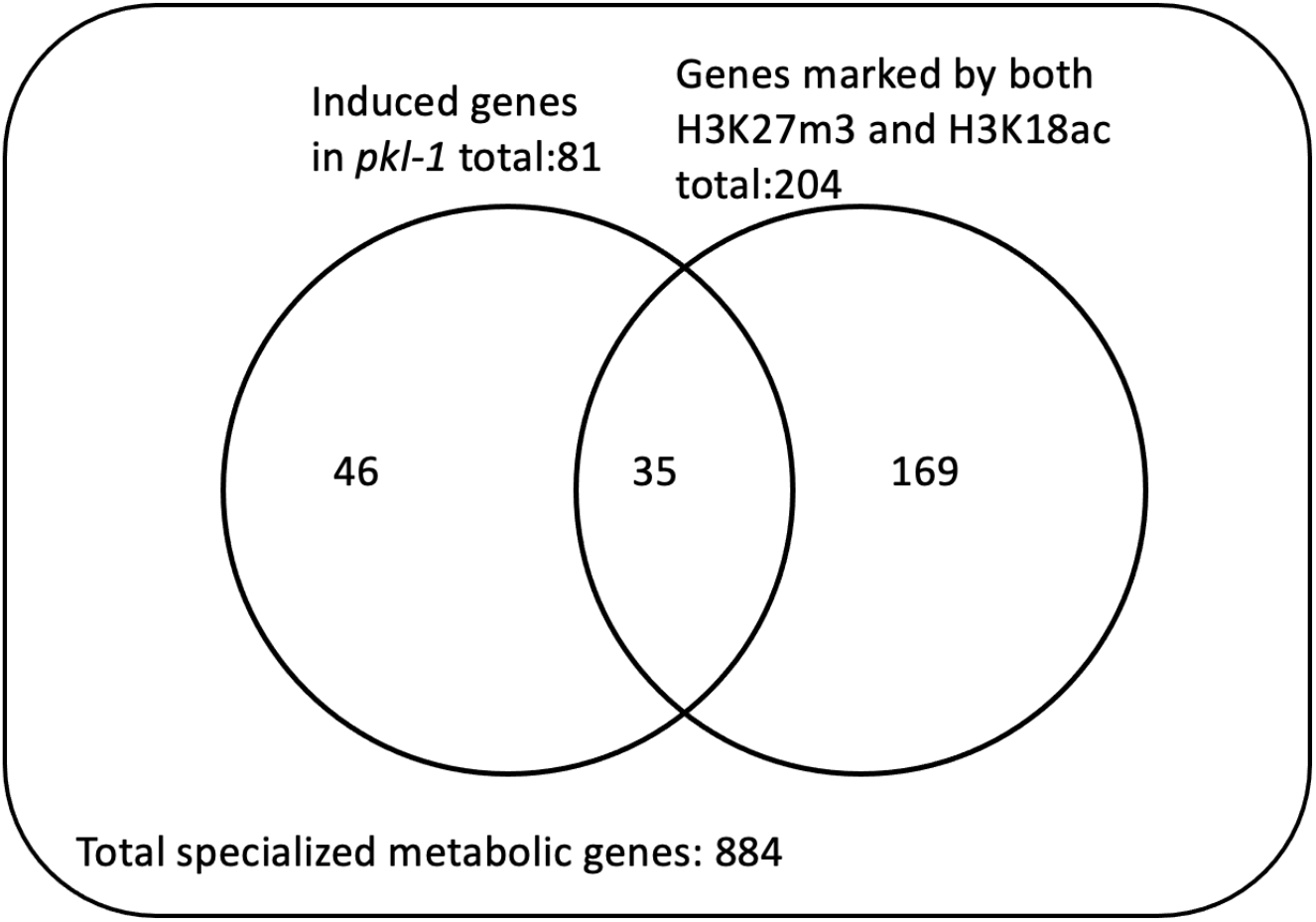
Among all the induced specialized metabolic genes in *pkl-1*, genes marked by both H3K27me3 and H3K18ac were more than expected by chance with hypergeometric test p value 1.8E-5.

**Fig S6.**
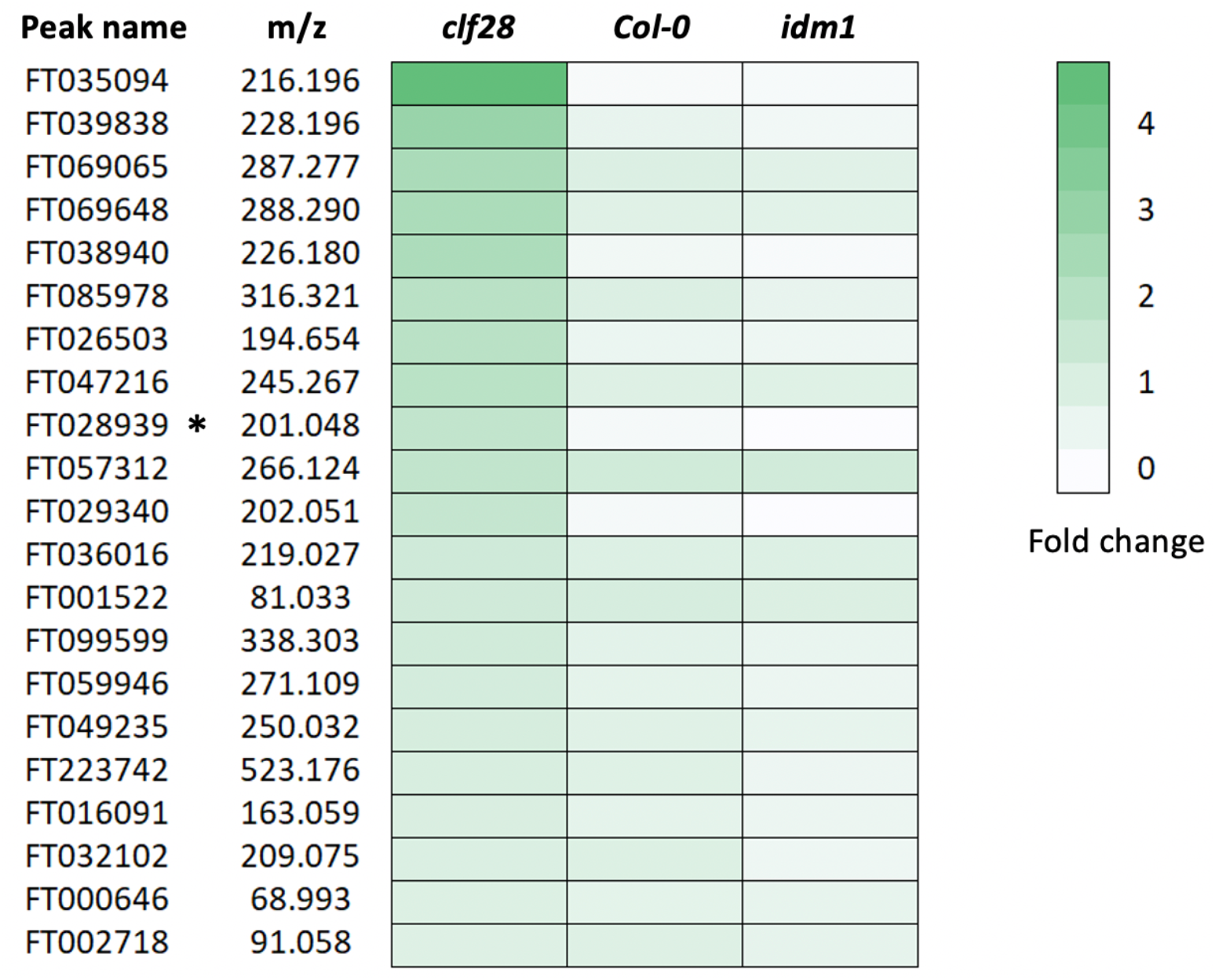
Heatmap of the 21 metabolites that accumulated the most in *clf28* plants and the least in *idm1* plants 3h after FLG22 treatment relative to water treatment. Significantly increased metabolites in response to 3h of FLG22 relative to water treatment in *clf28* plants were determined based on one-way ANOVA at p-value threshold of 0.05. Of the significantly induced compounds in *clf28*, only those with the highest accumulation level in *clf28* and the lowest in *idm1* plants were selected. The metabolites were ranked by the fold change in *clf28* plants. Each row represents one metabolite, labelled by its mass/charge ratio (m/z). Highlighted by an asterisk is camalexin (201.1468 m/z), which was validated by an authentic chemical standard.

**Table S2.**
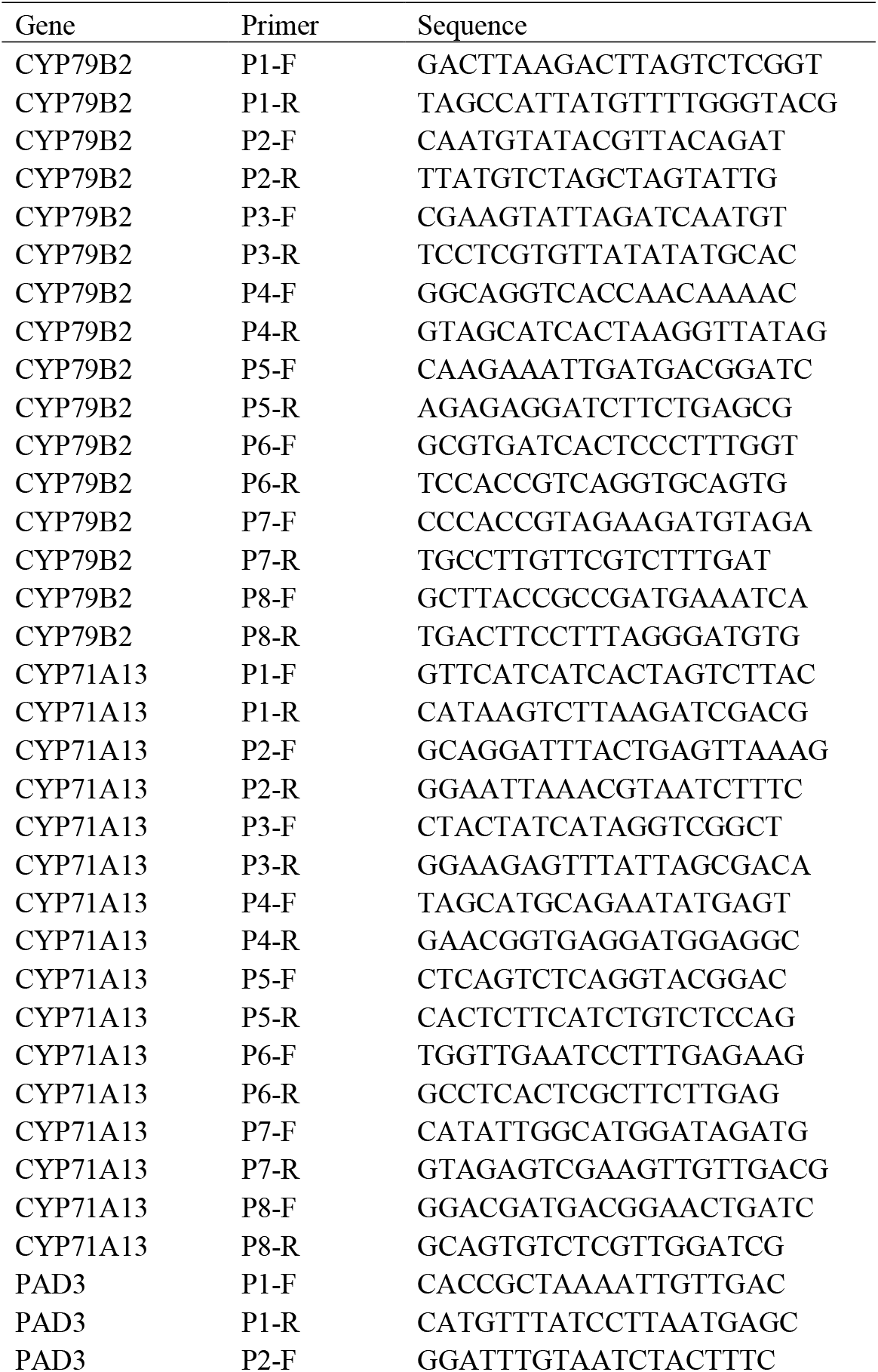
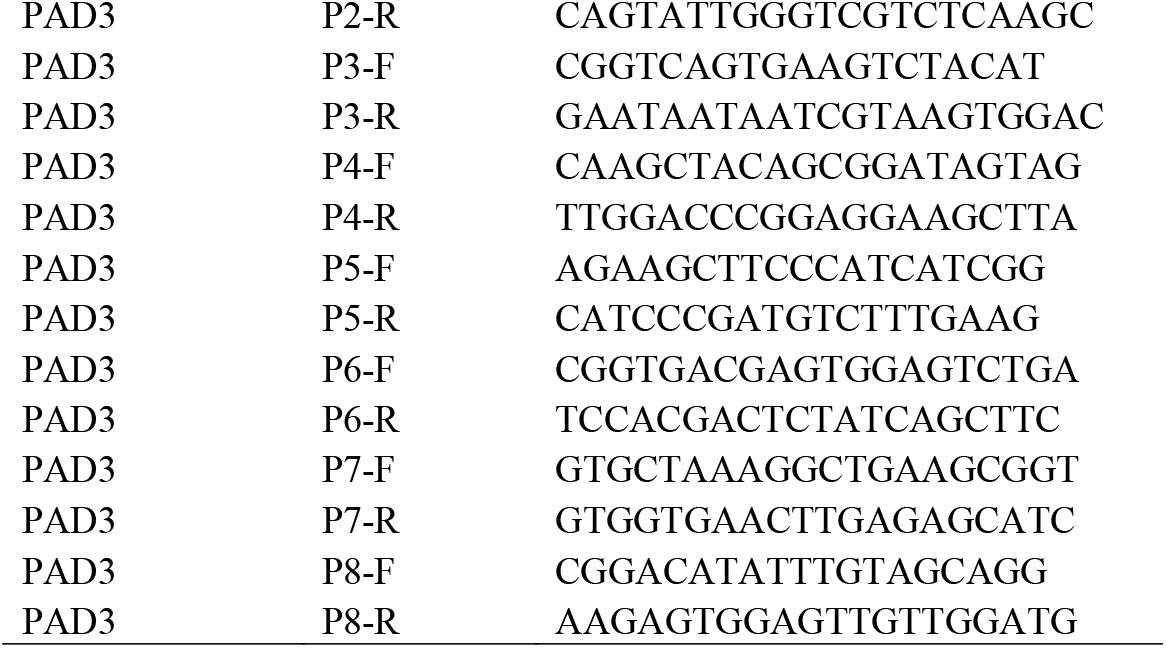
Primers used to quantify the abundance of H3K27me3 and H3K18ac in the genomic regions of camalexin biosynthesis genes using ChIP-qPCR in wild type and mutant plants with or without FLG22 treatment.

**Table S3.**
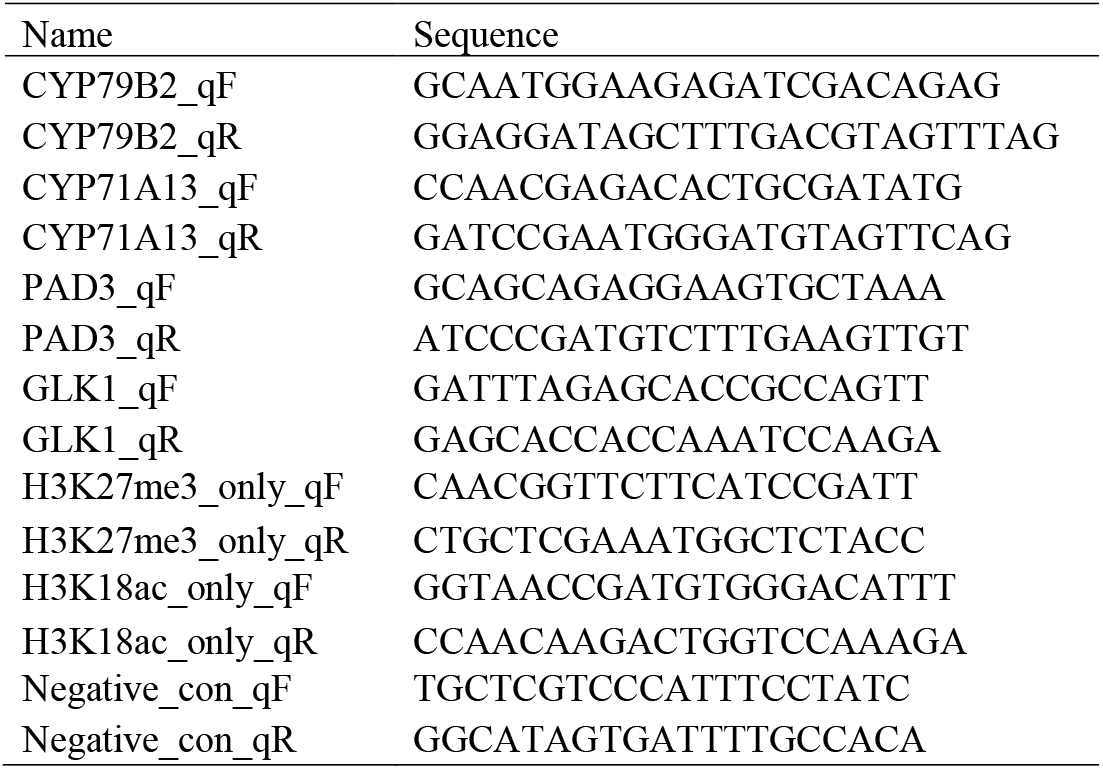
Primers used in the sequential ChIP-qPCR experiment to examine the co-localization of H3K27me3 and H3K18ac within camalexin biosynthesis genes.

**Table S4.**
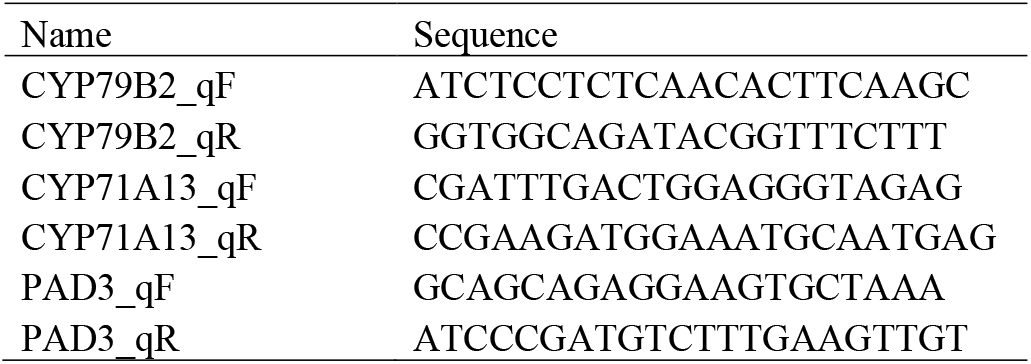
Primers used to examine the expression of camalexin biosynthesis genes under FLG22 induction using qPCR.

**Table S5.** Transcriptional kinetics of camalexin biosynthesis genes in wild type and mutant plants at different time points after FLG22 treatment. Fold change (FC) represents the expression level of these genes in samples treated with 1uM FLG22 for different amounts of time relative to their corresponding control tissues treated by water. Asterisks represent significantly increased gene expression levels at each time point based on two-tailed Student’s t test p value 0.05. Two-week old seedlings were used in this experiment with three biological replicates per experiment. The transcriptional kinetics assays were performed twice and showed similar results. sd represents standard deviation.

| Gene | Mutant line | 5m FC | 30m FC | 1h FC | 3h FC | 6h FC | 5m sd | 30m sd | 1h sd | 3h sd | 6h sd |
| --- | --- | --- | --- | --- | --- | --- | --- | --- | --- | --- | --- |
| *CYP71A13* | *Col-0* | 1.020 | 1.560* | 1.684* | 2.589* | 5.554* | 0.105 | 0.017 | 0.365 | 0.303 | 1.427 |
| *CYP71A13* | *pkl-1* | 1.773* | 2.429* | 2.418* | 3.743* | 9.452* | 0.089 | 0.279 | 0.560 | 0.551 | 1.615 |
| *CYP71A13* | *clf28* | 2.230* | 1.994* | 2.773* | 3.906* | 8.622* | 0.461 | 0.349 | 0.328 | 0.537 | 1.702 |
| *CYP71A13* | *idm1* | 1.164 | 1.100 | 1.903* | 1.554* | 2.857* | 0.029 | 0.119 | 0.007 | 0.095 | 0.250 |
| *CYP71A13* | *idm2* | 1.295 | 1.139 | 1.948* | 1.452* | 4.040* | 0.279 | 0.216 | 0.113 | 0.121 | 0.185 |
| *CYP71B2* | *Col-0* | 0.984 | 1.397 | 1.569* | 2.242* | 2.244* | 0.025 | 0.131 | 0.123 | 0.159 | 0.152 |
| *CYP71B2* | *pkl-1* | 1.781* | 1.847* | 2.186* | 3.266* | 3.497* | 0.387 | 0.441 | 0.084 | 0.589 | 0.612 |
| *CYP71B2* | *clf28* | 1.914* | 1.942* | 2.104* | 2.256* | 2.693* | 0.190 | 0.133 | 0.246 | 0.361 | 0.356 |
| *CYP71B2* | *idm1* | 0.920 | 1.016 | 1.080 | 1.745* | 1.539* | 0.278 | 0.240 | 0.381 | 0.397 | 0.127 |
| *CYP71B2* | *idm2* | 1.061 | 1.561* | 1.056 | 1.570* | 1.528* | 0.306 | 0.118 | 0.177 | 0.238 | 0.069 |
| *PAD3* | *Col-0* | 1.215 | 1.743* | 2.449* | 2.548* | 5.570* | 0.227 | 0.374 | 0.483 | 0.309 | 0.752 |
| *PAD3* | *pkl-1* | 1.559* | 2.283* | 3.111* | 3.905* | 7.058* | 0.142 | 0.075 | 0.012 | 0.372 | 0.512 |
| *PAD3* | *clf28* | 1.637* | 2.037* | 3.004* | 3.301* | 6.490* | 0.327 | 0.070 | 0.240 | 0.279 | 0.626 |
| *PAD3* | *idm1* | 1.017 | 1.282 | 1.763* | 2.191* | 2.901* | 0.028 | 0.175 | 0.087 | 0.245 | 0.368 |
| *PAD3* | *idm2* | 0.966 | 0.977 | 1.745* | 1.525* | 2.842* | 0.160 | 0.183 | 0.282 | 0.222 | 0.353 |

**Table S6.** Camalexin accumulation in wild type and mutant plants at different time points after FLG22 treatment. Con-mean represents the average metabolite content in control tissues treated with water. FLG-mean represents the average metabolite content in tissues treated with 1uM FLG22. Asterisks represent significantly increased camalexin content in FLG22 treated samples compared to their corresponding controls based on two-tailed Student’s t test p value 0.05. sd represents standard deviation. Two-week old seedlings were used in this experiment with three biological replicates per experiment. Camalexin content was measured twice and showed similar results.

| Time point | Line | Con-mean (nmol/g) | Con-sd | FLG-mean (nmol/g) | FLG-sd | Fold Change |
| --- | --- | --- | --- | --- | --- | --- |
| 5m | *col* | 0.242 | 0.032 | 0.278 | 0.009 | 1.148 |
| 5m | *pkl-1* | 0.311 | 0.072 | 0.28 | 0.083 | 0.898 |
| 5m | *clf28* | 0.235 | 0.015 | 0.247 | 0.031 | 1.049 |
| 5m | *idm1* | 0.23 | 0.058 | 0.253 | 0.047 | 1.106 |
| 5m | *idm2* | 0.275 | 0.051 | 0.336 | 0.071 | 1.224 |
| 30m | *col* | 0.305 | 0.021 | 0.316 | 0.054 | 1.038 |
| 30m | *pkl-1* | 0.319 | 0.05 | 0.255 | 0.07 | 0.801 |
| 30m | *clf28* | 0.304 | 0.01 | 0.321 | 0.02 | 1.152 |
| 30m | *idm1* | 0.23 | 0.009 | 0.228 | 0.018 | 0.992 |
| 30m | *idm2* | 0.236 | 0.064 | 0.255 | 0.036 | 1.082 |
| 1h | *col* | 0.226 | 0.039 | 0.279 | 0.058 | 1.233 |
| 1h | *pkl-1* | 0.214 | 0.017 | 0.287 | 0.044 | 1.641 |
| 1h | *clf28* | 0.144 | 0.031 | 0.288 | 0.022 | 1.999* |
| 1h | *idm1* | 0.255 | 0.098 | 0.208 | 0.116 | 0.896 |
| 1h | *idm2* | 0.175 | 0.016 | 0.243 | 0.024 | 1.388 |
| 3h | *Col* | 0.214 | 0.059 | 0.339 | 0.021 | 1.580* |
| 3h | *pkl-1* | 0.189 | 0.041 | 0.467 | 0.017 | 2.587* |
| 3h | *clf28* | 0.162 | 0.008 | 0.495 | 0.02 | 3.162* |
| 3h | *idm1* | 0.202 | 0.05 | 0.245 | 0.033 | 1.361 |
| 3h | *idm2* | 0.219 | 0.035 | 0.268 | 0.059 | 1.223 |
| 6h | *Col* | 0.162 | 0.017 | 0.409 | 0.055 | 2.548* |
| 6h | *pkl-1* | 0.183 | 0.01 | 0.482 | 0.025 | 2.633* |
| 6h | *clf28* | 0.152 | 0.005 | 0.433 | 0.013 | 2.545* |
| 6h | *idm1* | 0.18 | 0.043 | 0.273 | 0.052 | 1.867* |
| 6h | *idm2* | 0.147 | 0.03 | 0.238 | 0.018 | 1.613* |

## References

1. Barco B, Clay NK. Hierarchical and Dynamic Regulation of Defense-Responsive Specialized Metabolism by WRKY and MYB Transcription Factors. Front Plant Sci 10, 1775 (2019).

2. Grotewold E. Plant metabolic diversity: a regulatory perspective. Trends Plant Sci 10, 57–62 (2005).

3. Martin C, Ellis N, Rook F. Do transcription factors play special roles in adaptive variation? Plant Physiol 154, 506–511 (2010).

4. Stotz HU, et al. Role of camalexin, indole glucosinolates, and side chain modification of glucosinolate-derived isothiocyanates in defense of Arabidopsis against Sclerotinia sclerotiorum. Plant J 67, 81–93 (2011).

5. Klein AP, Anarat-Cappillino G, Sattely ES. Minimum set of cytochromes P450 for reconstituting the biosynthesis of camalexin, a major Arabidopsis antibiotic. Angew Chem Int Ed Engl 52, 13625–13628 (2013).

6. Schuhegger R, et al. CYP71B15 (PAD3) catalyzes the final step in camalexin biosynthesis. Plant Physiol 141, 1248–1254 (2006).

7. Denoux C, et al. Activation of defense response pathways by OGs and Flg22 elicitors in Arabidopsis seedlings. Mol Plant 1, 423–445 (2008).

8. Frerigmann H, et al. Regulation of Pathogen-Triggered Tryptophan Metabolism in Arabidopsis thaliana by MYB Transcription Factors and Indole Glucosinolate Conversion Products. Mol Plant 9, 682–695 (2016).

9. Stahl E, et al. Regulatory and Functional Aspects of Indolic Metabolism in Plant Systemic Acquired Resistance. Mol Plant 9, 662–681 (2016).

10. Zhou J, et al. Differential Phosphorylation of the Transcription Factor WRKY33 by the Protein Kinases CPK5/CPK6 and MPK3/MPK6 Cooperatively Regulates Camalexin Biosynthesis in Arabidopsis. Plant Cell 32, 2621–2638 (2020).

11. Birkenbihl RP, Diezel C, Somssich IE. Arabidopsis WRKY33 is a key transcriptional regulator of hormonal and metabolic responses toward Botrytis cinerea infection. Plant Physiol 159, 266–285 (2012).

12. Stricker SH, Koferle A, Beck S. From profiles to function in epigenomics. Nat Rev Genet 18, 51–66 (2017).

13. Vihervaara A, Duarte FM, Lis JT. Molecular mechanisms driving transcriptional stress responses. Nat Rev Genet 19, 385–397 (2018).

14. Jiang D, Berger F. Histone variants in plant transcriptional regulation. Biochim Biophys Acta Gene Regul Mech 1860, 123–130 (2017).

15. Henikoff S, Greally JM. Epigenetics, cellular memory and gene regulation. Curr Biol 26, R644–648 (2016).

16. Carter B, et al. The Chromatin Remodelers PKL and PIE1 Act in an Epigenetic Pathway That Determines H3K27me3 Homeostasis in Arabidopsis. Plant Cell 30, 1337–1352 (2018).

17. Aranda S, Mas G, Di Croce L. Regulation of gene transcription by Polycomb proteins. Sci Adv 1, e1500737 (2015).

18. Zhang X, Bernatavichute YV, Cokus S, Pellegrini M, Jacobsen SE. Genome-wide analysis of mono-, di- and trimethylation of histone H3 lysine 4 in Arabidopsis thaliana. Genome Biol 10, R62 (2009).

19. Lauberth SM, et al. H3K4me3 interactions with TAF3 regulate preinitiation complex assembly and selective gene activation. Cell 152, 1021–1036 (2013).

20. Bernstein BE, et al. A bivalent chromatin structure marks key developmental genes in embryonic stem cells. Cell 125, 315–326 (2006).

21. Voigt P, Tee W-W, Reinberg D. A double take on bivalent promoters. Genes Dev 27, 1318–1338 (2013).

22. Dattani A, et al. Epigenetic analyses of planarian stem cells demonstrate conservation of bivalent histone modifications in animal stem cells. Genome Res 28, 1543–1554 (2018).

23. Sachs M, Onodera C, Blaschke K, Ebata KT, Song JS, Ramalho-Santos M. Bivalent chromatin marks developmental regulatory genes in the mouse embryonic germline in vivo. Cell Rep 3, 1777–1784 (2013).

24. Jiang D, Wang Y, Wang Y, He Y. Repression of FLOWERING LOCUS C and FLOWERING LOCUS T by the Arabidopsis Polycomb repressive complex 2 components. PLoS One 3, e3404 (2008).

25. Zeng Z, Zhang W, Marand AP, Zhu B, Buell CR, Jiang J. Cold stress induces enhanced chromatin accessibility and bivalent histone modifications H3K4me3 and H3K27me3 of active genes in potato. Genome Biol 20, 123 (2019).

26. Luo C, Sidote DJ, Zhang Y, Kerstetter RA, Michael TP, Lam E. Integrative analysis of chromatin states in Arabidopsis identified potential regulatory mechanisms for natural antisense transcript production. Plant J 73, 77–90 (2013).

27. Zhang Q, et al. Asymmetric epigenome maps of subgenomes reveal imbalanced transcription and distinct evolutionary trends in Brassica napus. Mol Plant, (2020).

28. Wang C, et al. Genome-wide analysis of local chromatin packing in Arabidopsis thaliana. Genome Res 25, 246–256 (2015).

29. Roudier F, et al. Integrative epigenomic mapping defines four main chromatin states in Arabidopsis. EMBO J 30, 1928–1938 (2011).

30. Schlapfer P, et al. Genome-Wide Prediction of Metabolic Enzymes, Pathways, and Gene Clusters in Plants. Plant Physiol 173, 2041–2059 (2017).

31. Qian W, et al. Regulation of active DNA demethylation by an alpha-crystallin domain protein in Arabidopsis. Mol Cell 55, 361–371 (2014).

32. Qian W, et al. A histone acetyltransferase regulates active DNA demethylation in Arabidopsis. Science 336, 1445–1448 (2012).

33. Zipfel C, et al. Bacterial disease resistance in Arabidopsis through flagellin perception. Nature 428, 764–767 (2004).

34. Chen Z, Li S, Subramaniam S, Shyy JY, Chien S. Epigenetic Regulation: A New Frontier for Biomedical Engineers. Annu Rev Biomed Eng 19, 195–219 (2017).

35. Hou X, Zhou J, Liu C, Liu L, Shen L, Yu H. Nuclear factor Y-mediated H3K27me3 demethylation of the SOC1 locus orchestrates flowering responses of Arabidopsis. Nat Commun 5, 4601 (2014).

36. Chae L, Kim T, Nilo-Poyanco R, Rhee SY. Genomic signatures of specialized metabolism in plants. Science 344, 510–513 (2014).

37. Karp PD, Latendresse M, Caspi R. The pathway tools pathway prediction algorithm. Stand Genomic Sci 5, 424–429 (2011).

38. Zhang P, et al. Creation of a Genome-Wide Metabolic Pathway Database for Populus trichocarpa Using a New Approach for Reconstruction and Curation of Metabolic Pathways for Plants. 153, 1479–1491 (2010).

39. Sham PC, Purcell SM. Statistical power and significance testing in large-scale genetic studies. Nat Rev Genet 15, 335–346 (2014).

40. Warnes GR, et al. gplots: Various R programming tools for plotting data. 2, 1 (2009).

41. Wickham H. ggplot2: elegant graphics for data analysis. Springer (2016).

42. Zhu JY, Oh E, Wang T, Wang ZY. TOC1-PIF4 interaction mediates the circadian gating of thermoresponsive growth in Arabidopsis. Nat Commun 7, 13692 (2016).

43. Saleh A, Alvarez-Venegas R, Avramova Z. An efficient chromatin immunoprecipitation (ChIP) protocol for studying histone modifications in Arabidopsis plants. Nat Protoc 3, 1018–1025 (2008).

44. Love MI, Huber W, Anders S. Moderated estimation of fold change and dispersion for RNA-seq data with DESeq2. Genome Biol 15, 550 (2014).

45. Smith CA, Want EJ, O’Maille G, Abagyan R, Siuzdak G. XCMS: processing mass spectrometry data for metabolite profiling using nonlinear peak alignment, matching, and identification. Anal Chem 78, 779–787 (2006).

